# The hidden secrets of the dental calculus: Calibration of a mass spectrometry protocol for dental calculus protein analysis

**DOI:** 10.1101/2022.07.27.501661

**Authors:** Omer Bender, Dana Tzabag, Talia Gavish, Noa Meyrom, Neta Zamir, Hila May, Rachel Sarig, Daniel Z. Bar

## Abstract

Dental calculus is a solid deposit that forms and accumulates on the tooth surface, entrapping oral microorganisms, biomolecules, and other micro-debris found in the oral cavity. Mass spectrometry analysis of its protein content opens a vista into the subject’s diet, oral flora, and even some aspects of health, thus providing new insight and expanding our knowledge of archaic cultures. Multiple experimental protocols have been proposed for the optimal extraction of proteins from dental calculus. Herein, we compared various experimental conditions in order to calibrate and validate a protocol for protein extraction. Our results show that a high concentration of acetic acid followed by mechanical crushing and sonication provided the highest protein yield, while acetone precipitation enabled the identification of more distinct proteins. We validated this protocol using archeological samples, identifying human and microbial proteins in specimens from the 8th and 17th centuries (approximately 250–1300 years ago). These findings demonstrate that the developed protocol is useful for studying excavated archaeological samples and that it might be utilized to explore the biohistory, dietary habits, and microbiome of archaic populations.

## Introduction

Dental calculus, also called tooth tartar, is a solid deposit that forms and accumulates on the tooth surfaces and creates a firm bond with the teeth that can only be removed by mechanical means. Dental calculus originates from a mineralized oral bacterial plaque that naturally forms on tooth surfaces where there is a constant supply of saliva. The mineralization process is caused by the deposition of calcium phosphate crystals from the saliva into the bacterial plaque ^1^. Since calculus formation is related to saliva sources, it is commonly seen over the buccal surfaces of maxillary molars and lingual surfaces of mandibular anterior teeth, where the salivary ducts open into the oral cavity ^2^.

The calcification process of dental plaque is dynamic: it begins with the adsorption of salivary proteins on the enamel surfaces, followed by adherence of a structurally organized biofilm, which eventually calcifies in several phases of mineralization. During the process, the precipitation of mineral ions (calcium and phosphate) from the saliva and gingival crevicular fluid is settled in the bacterial biofilm, together with additional oral microorganisms, biomolecules, and other micro-debris sourced directly from the human mouth ^3^.

The calculus matrix consists mainly of inorganic salts and mineral ions deriving from saliva (e.g., hydroxyapatite, calcium, whitlockite, and phosphorus). It also consists of organic compounds, including microfossils (e.g., starch granules, phytoliths, and plant metabolites) and biomolecules (e.g., proteins, lipids, and nucleic acids) derived from consumed foods, the oral microbiome, or the host ^3,4^.

As dental calculus captures and preserves microparticles throughout an individual’s lifetime, it may contain valuable information. Due to its densely mineralized nature, calculus can serve as a long-term abundant reservoir of ancient proteins that can last for extended time periods and preserve numerous biomolecules containing valuable information. Therefore, archaeological dental calculus serves as a fossilised record of the oral environment. A single sample of calculus contains numerous molecules that can provide information about the individual’s genome ^5,6^, oral microbiome and health ^7^, diet ^8,9^, and even occupational activities ^10^.

Changes in prehistoric populations’ lifestyles are determined by diet, culture, and habits ^11^ therefore, these parameters can be reflected by the composition of dental calculus deposits ^12^. Accordingly, analysis of dental calculus offers a unique opportunity to reveal the everyday function of the individuals and gain insight into the lifestyle and dietary habits of prehistoric populations. In addition to other dietary molecules, the dental calculus specifically preserves food-related proteins, and by a proper extraction method, we can detect proteomic evidence of a variety of food ingredients ^13^.

Moreover, as the human mouth is a habitat ecosystem for a variety of microorganisms, the dental calculus preserves proteins associated with the oral microbiota and might contain disease-associated microorganisms, including viruses and fungi ^14,15^. Archaeological dental calculus is a rich source of ancient proteins derived from the microbiome and thus serves as a tool for palaeomicrobiology (the study of microorganisms from ancient sources) and has a great potential to elucidate the evolution of the human microbiome. This knowledge might shed light on human-associated microbes in a variety of acute and chronic diseases and provide a valuable and vital source of information for assessing individual health ^14,16^.

### Mass spectrometry (MS) for quantitative proteomics analyses

MS is a powerful method for hypothesis-free and hypothesis-driven identification of proteins. By applying this method to mineral deposits, we can detect the presence of protein molecules in the sample. The MS-based proteomics workflow requires an extraction procedure for the existing proteins, which are then digested into peptides. The peptide mixture is eventually analysed by MS ^8,17^. The timeframe in which these data can be retrieved varies, and depends, among others, on the preservation conditions of the archaeological specimens in the excavated sites.

### Limitations

In recent years, dental calculus has become the subject of an increasing number of investigations that concern different research fields in biology ^5–9,12,14,17–19^. However, the quantity of archaeological dental calculus available per individual is generally low, sometimes only tens of milligrams per sample. This is partially due to preservation protocols that historically included cleaning of the archaeological specimens and removal of any residual calculus, or, in cases that only few isolated teeth were recovered from the sites. This factor limits the number of analyses that can be performed for the extraction of biomolecules from different sources ^5^. According to these limitations, it is evident that the amount of dental calculus has become the main limiting factor for obtaining data. Thus, a reliable, reproducible and effective protein extraction protocol allowing to obtain considerable information from archaeological samples is highly required.

Recent studies suggested various extraction protocols to evaluate archaic dental calculus using metaproteomics ^17,19^, yet, there is still a debate as for what is the most effective protocol for such purposes. Herein, we provide an optimised protocol for protein content analysis from dental calculus samples. We compared various conditions for decalcification, mechanical crushing, sonication, protein precipitation and MS gradient times. These resulted in a single optimized protocol (Supplementary File 1) that can be used to deduce individual dietary practices and inform us on oral health and oral microbiome composition in a specific population.

## Material and Methods

For MS calibration, a minimum amount of 10-15 mg of dental calculus was considered necessary to obtain significant results in each experiment. For accurate comparison and for optimal protocol establishment, all dental calculus used was collected from 6 patients at the clinic following signed and informed consent. Medical confidentiality was guaranteed regarding the identity of patients. The study was approved by the ethical committee of Tel Aviv University (Number 0005110-2). The fresh dental calculus was obtained using a sterile dental curette and stored in sterile 2.0 mL tubes until further analysis.

The established protocol was further applied upon the archaeological samples retrieved from the anthropological collection at the Dan David Center for Human Evolution and Biohistory Research, Sackler Faculty of Medicine, Tel Aviv University. Two samples were obtained from excavated sites dated to Late Arab (the 17the century) and Early Arabs (8th century).

### Sample Preparation

In order to analyse the proteins captured in human calculus, dental calculus was decalcified with 15%/20%/25% acetic acid overnight, the pellet was treated with\without mechanical sonication and sonication. Then, prior to MS analysis, samples were cleaned with different precipitation methods to remove detergents followed by 1 or 2 hours run time for MS analysis (**Fig. 1**). All the steps are detailed below.

**Figure 1:**
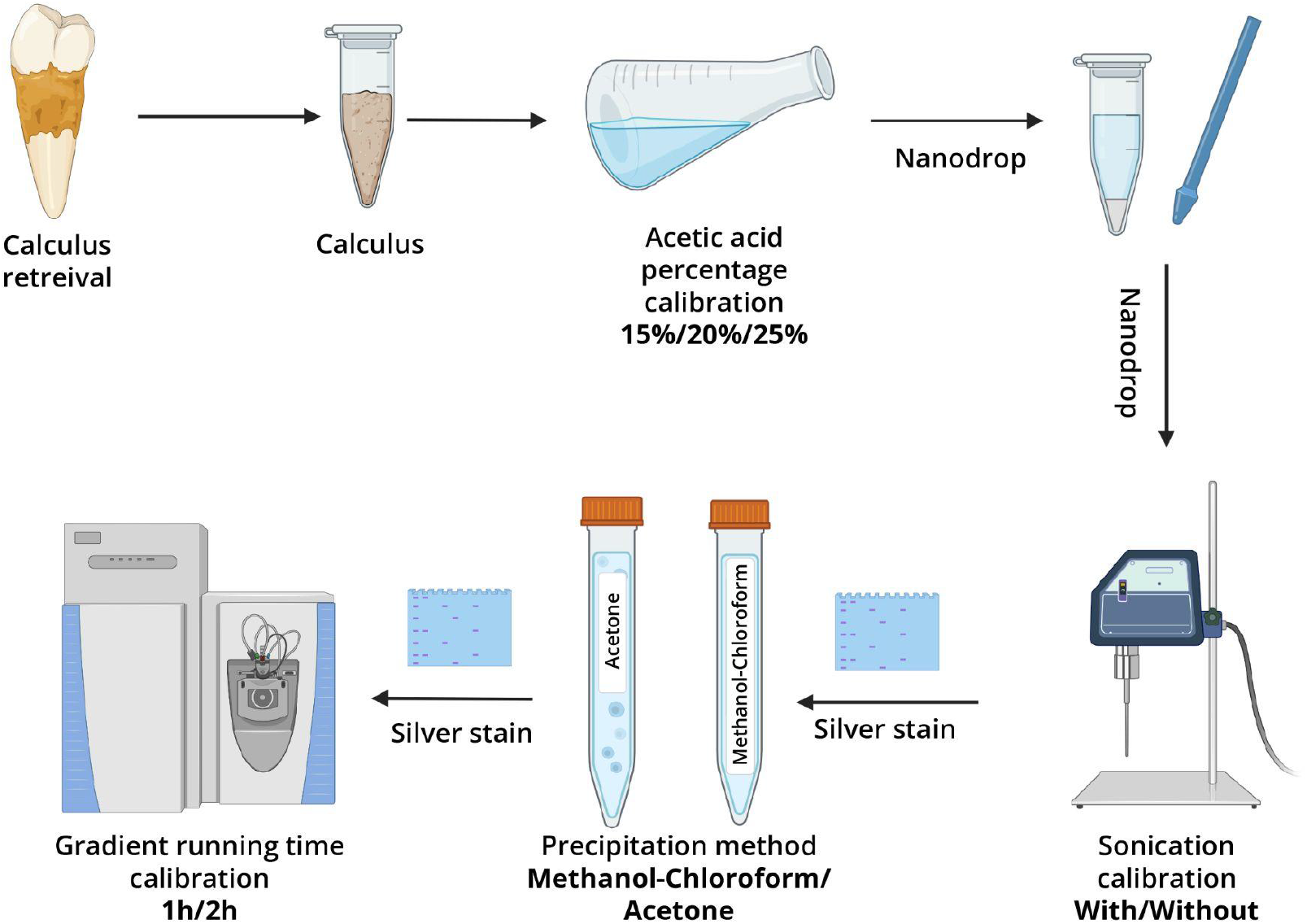
Protocol schematic overview.

### Decalcification

Upon usage, the fresh dental calculus samples retrieved from patients were washed 3 times with PBS buffer for 10 minutes. After each wash cycle, the calculus was spun down at 10,000 RCF. The calculus samples were then demineralized in 15%, 20% or 25% acetic acid (Biolab, Israel) overnight on a 3D orbital shaker. Following demineralization, samples were grounded mechanically using an Eppendorf micro pastel (Merck). Samples were washed 4 times with PBS for 10 minutes. After each cycle of washing, samples were spun down at 10,000 RCF. Calculus samples were resuspended in lysis buffer (2M Guanidine Hydrochloride (Merck), 10mM Chloroacetamide (Bar Naor - Israel) and 5mM HEPES (Merck), pH 7.5), followed by sonication for 25 minutes at 40%. Samples were spun down at 21,000 RCF for 30 minutes at 4°C.

### Residual Protein Loss during Decalcification

Protein concentrations were measured using the A205 preprogrammed direct absorbance application of Nanodrop One C (Thermo Fisher Scientific). Measurements were performed following acetic acid incubation and after the mechanical crushing to exclude the possibility of protein loss in washes.

### Protein Precipitation

Following optional sonication, each sample was divided into 2 equal aliquots which were further processed using two different precipitation methods:

#### 1. Methanol-Chloroform

Methanol at 4 times (600μl) the sample volume (150μl) was added and vortexed thoroughly. Next, chloroform at the original sample volume was added and vortexed. Next, 400 μl of double distilled water (DDW) was added and the sample was vortexed thoroughly until the mixture became cloudy with precipitation. The sample was centrifuged at 14,000 RCF for 1 minute. Three layers were apparent: top aqueous layer, circular flake of protein. The top layer was removed without disturbing the protein at the interface. Methanol at 4 times of sample volume was added and vortexed. The sample was centrifuged at 20,000 RCF for 5 minutes, and excess methanol was discarded. Then, the sample was placed under vacuum and resuspended with 100 uL of 100 mM HEPES buffer (Merck).

#### 2. Acetone

Cold (−20°C) acetone at 4 times of sample volume was added and vortexed thoroughly. The sample was then incubated at -80°C for 60 minutes and centrifuged at 15,000 RCF for 10 minutes at 4°C. The supernatant was discarded. Then, the sample was placed under vacuum for complete removal of the acetone and resuspended with 100 uL of 100 mM HEPES buffer (Merck).

### Protein Processing

Following protein precipitation, samples underwent reduction by adding Dithiothreitol (Merck) to a final concentration of 10mM for 60 min at 55°C, followed by alkylation by adding Iodoacetamide (Merck) to a final concentration of 18.75 mM for 30 minutes at room temperature. Subsequently, samples were digested overnight at 37°C with 10 nG MS Grade Trypsin (Promega). Trypsinization was quenched with 1 μL of 0.1% Trifluoroacetic acid (TFA) (ThermoFisher).

### Solid Phase Extraction Stage Tips for Detergent Removal

Prior to mass spectrometry, samples were cleaned with a solid phase extraction method using 2 layers of C18 Empore™ SPE Disks (Merck) followed by 2 layers of SCX strong cation exchange Empore™ SPE Disks (Merck). Samples were then lyophilized.

### MS analysis

MS analysis was performed at the National Proteomics Unit at the Technion, Israel. The peptides were resolved by reverse-phase chromatography on 0.075 × 300-mm fused silica capillaries packed with Reprosil reversed phase material (Dr Maisch GmbH, Germany). The peptides were eluted using a linear 60 or 120 minutes gradient of 5 to 28%, a 15 minutes gradient of 28 to 95%, and 15 minutes at 95% Acetonitrile with 0.1% formic acid in water at flow rates of 0.15 μl/min. MS analysis was performed by Q Exactive HF mass spectrometer (Thermo) in a positive mode (m/z 300–1800, resolution 120,000 for MS1 and 15,000 for MS2) using repetitively full MS scan followed by high collision dissociation (HCD, at 27 normalized collision energy) of the 20 most dominant ions (≥1 charges) selected from the first MS scan. The AGC settings were 3×10 6 for the full MS and 1×10 5 for the MS/MS scans. The intensity threshold for triggering MS/MS analysis was 1.3×10 5. A dynamic exclusion list was enabled with exclusion duration of 20s.

### Silver Stain

15% SDS-PAGE gels were prepared in advance. Gels were made with 2 parts. Lower and stacking gel. Lower gel was made with 1.2 mL DDW, 5mL 30% Acrylamide 29:1 (Bar-Naor, Israel), 1.25 mL 1.5M Tris-HCl pH 8.8, 50 uL 10% Sodium Dodecyl Sulfate (SDS, Thermo-Fisher), 25 uL 10% Ammonium Persulfate (APS, Merck), 2.5 uL TEMED (Bar-Naor, Israel). Stacking Gel was made with with 2.3 mL DDW, 500 uL 30% Acrylamide 29:1, 950 uL 0.5M Tris-HCl pH 8.8, 37.5 uL 10% SDS, 18.75 uL 10% APS, 3.75 uL TEMED.

Samples were loaded with X4 Laemli Sample Buffer (LSB, Bio-Rad) and put in a preheated heating block set to 95°C for 5 minutes. Samples were then separated on the 15% SDS-PAGE gels with 120 mV constant voltage. Gels were then stained using MS compatible silverquest staining kit (Thermo - Fisher LC6070) according to manufacturer instructions.

### Archeological dental calculus samples

Following the calibration of the protocol, the procedure was applied to the dental calculus samples retrieved from the archeological sites. Each sample was treated as stated in our protocol.

### Bioinformatic analysis

To assess the effect of precipitation method and gradient time on the obtained data, we searched the data for peptides and proteins from two sources, expected to be present in our sample: proteins of human origin and from *E. Coli*. Protein raw data files were run on the MaxQuant V2.1.0.0 software (Max Planck institute). Protein minimal peptide count per protein was set to 3, with an FDR set to 10% per peptide.The following protein FASTA files were used:

- Homo Sapiens (Human) proteome ID UP000005640.
- Escherichia Coli - proteome ID UP000036496.

## Results

### Decalcification optimization

To enable efficient extraction of proteins from the sample, the dental calculus first needed to be decalcified. This is traditionally done by a combination of acid treatment with mechanical disintegration of the sample. Here we used different concentrations of acetic acid (AA, 15%, 20%, 25%). Samples were decalcified in the AA followed by manual crushing of the calculus using a micro-pestle. Next, we tested the effect of sonication on protein release. Spectrophotometric measurements confirmed no protein loss from the decalcification and mechanical crushing.

Protein yield increased with acid concentration and required mechanical crushing (**Fig. 2**). Sonication had a positive but very mild effect on protein yield. The greatest amount of protein yield was achieved using 25% AA on followed by mechanical crushing of the calculus and sonication. However, protein elution was not complete, and repetitive elution may be advised for limited samples.

**Figure 2:**
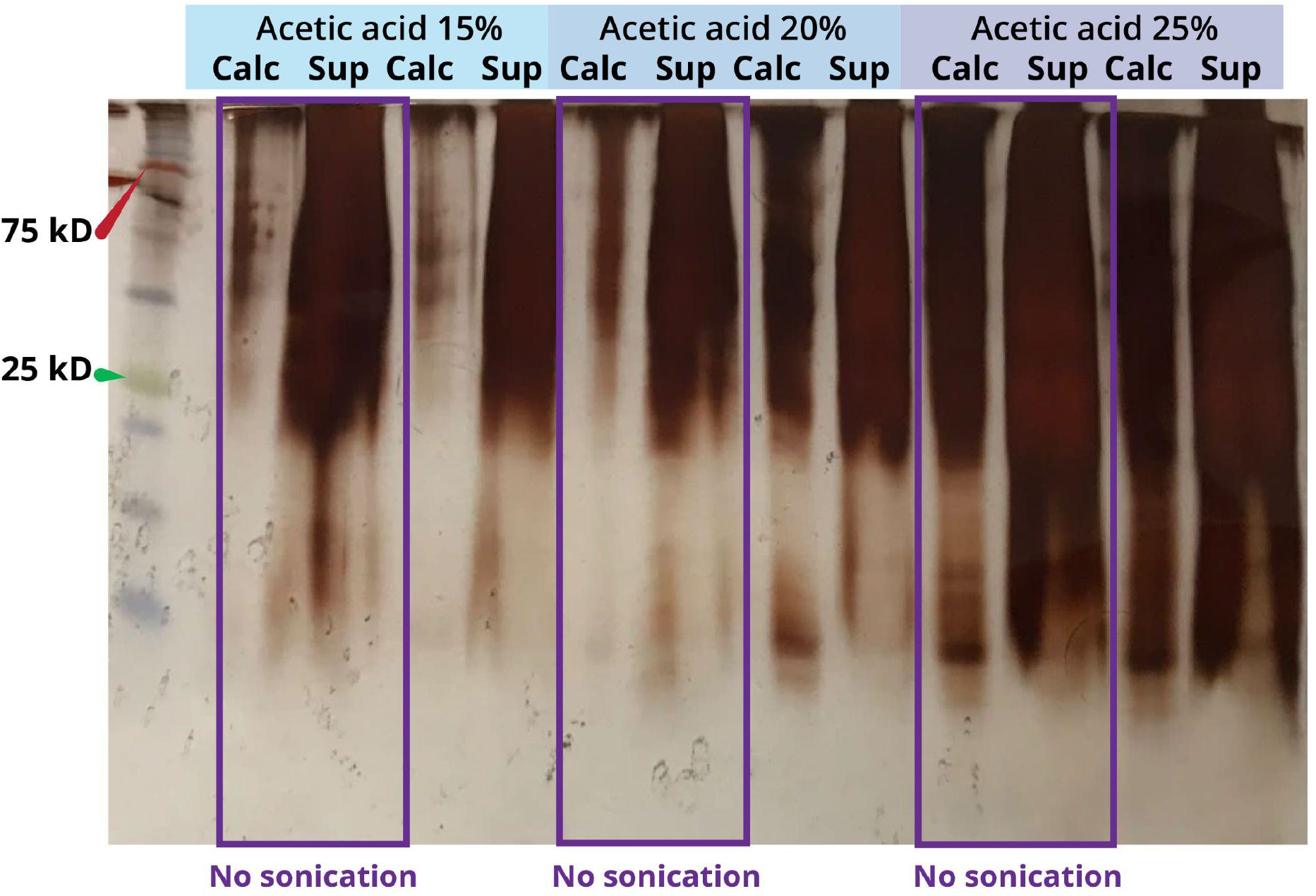
Silver-staining estimate of protein yield under different conditions. Calc. - calculus. Sup. - supernatant.

### Protein precipitation

Prior to protein digestion and mass spectrometry analysis, the protein content should be purified. We have tested which protein precipitation method results in higher protein yield. Without protein purification, no protein digestion using trypsin was detected (**Fig. 3**). Both methanol-chloroform and acetone precipitation methods facilitated sample purification to allow the full cleavage of the proteins into peptides (**Fig. 3**). However, the protein yield retrieved from acetone precipitation was lower compared to methanol-chloroform precipitation.

**Figure 3:**
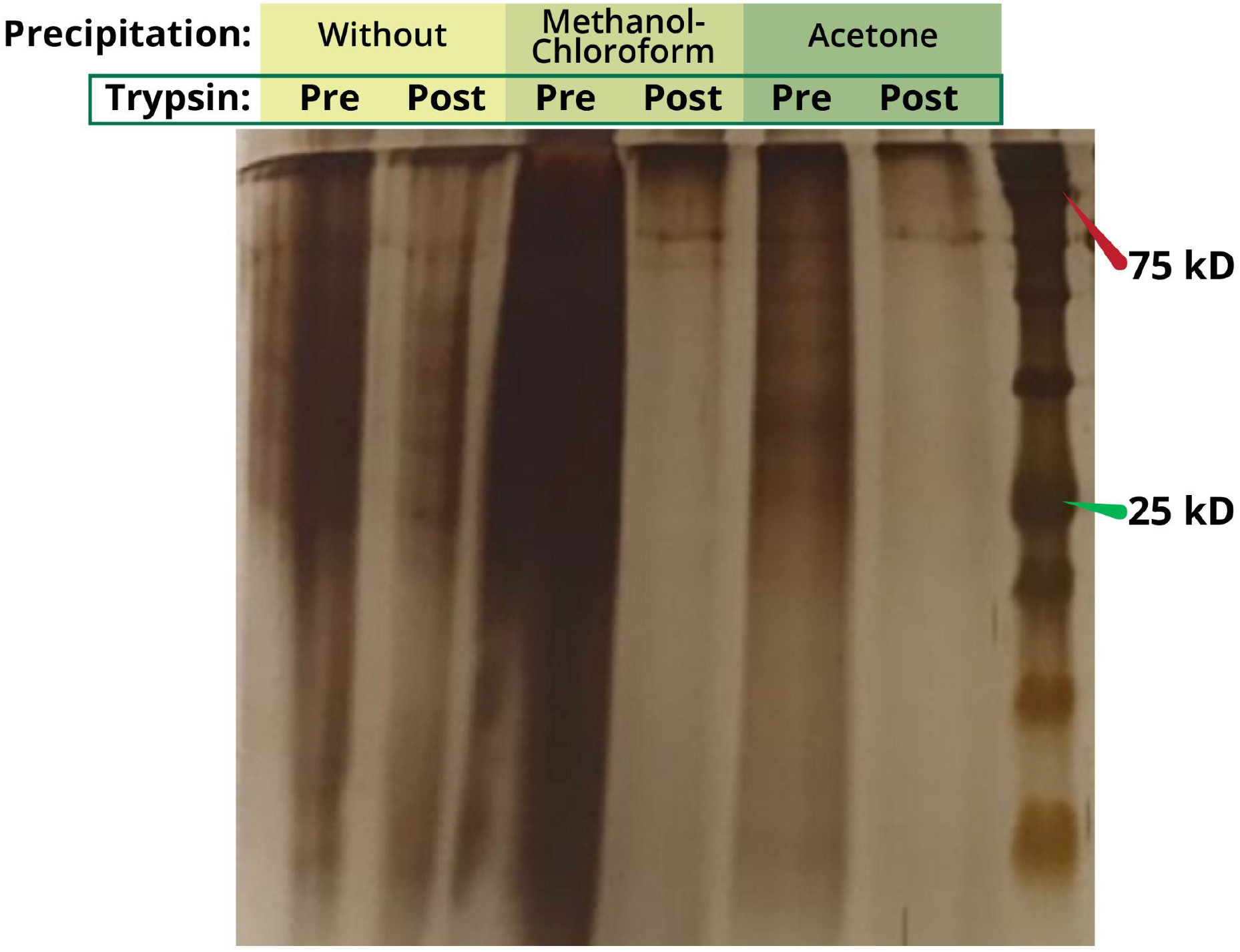
Silver-staining of proteins separated on an SDS-PAGE gel following either of two methods of precipitation. Methanol-chloroform and acetone precipitation products before and after trypsinization of the protein samples are shown.

### Gradient time optimization

Protein MS typically relies on peptide elution along an organic solvent gradient. Here, we compared 1-hour vs. 2-hour time gradients of protein samples following two different precipitation methods: methanol-chloroform and acetone (**Fig. 4 A-D**).

**Figure 4:**
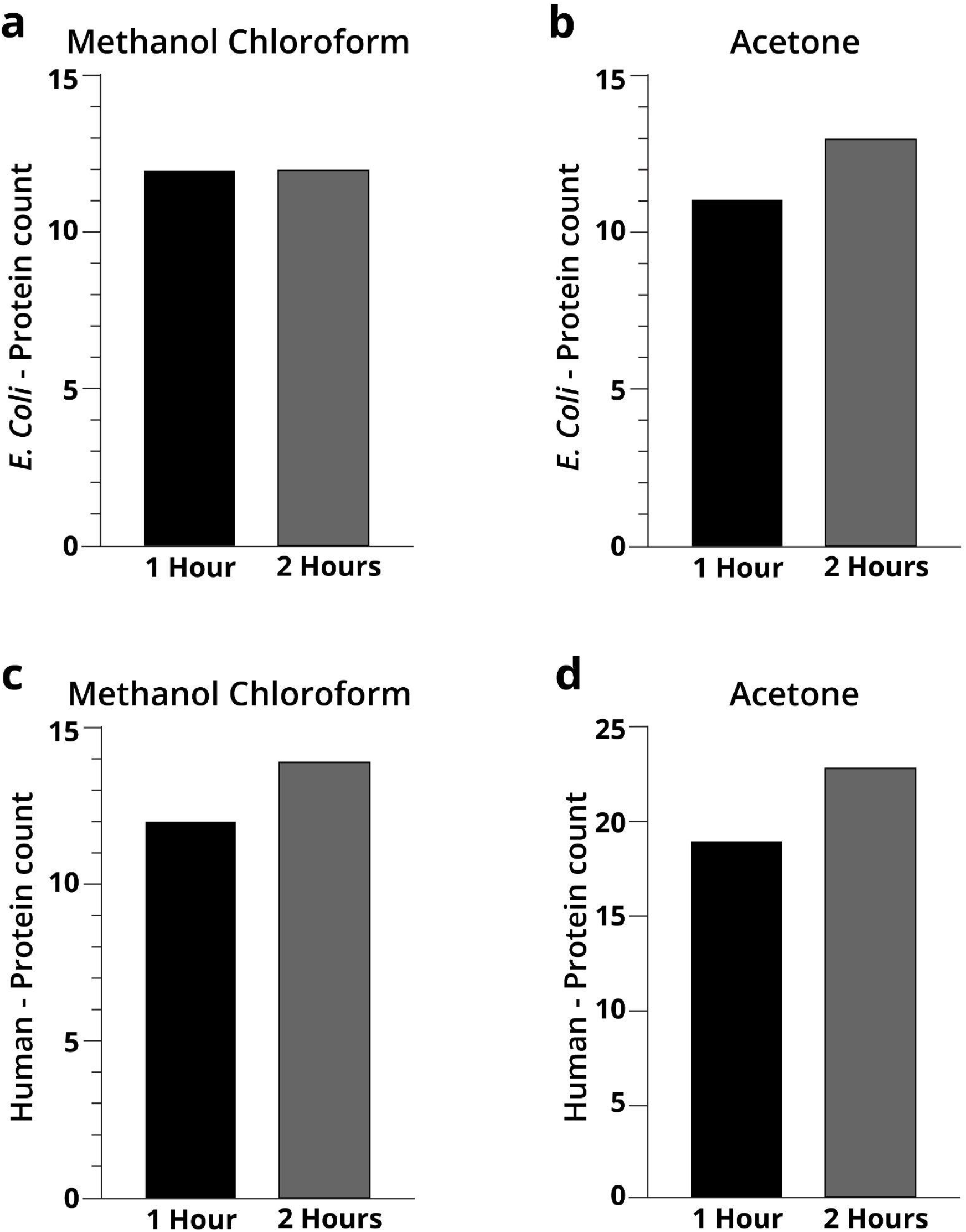
Comparison of the MS samples running time. **A**. 1 vs 2 hours run of *E. Coli* proteins after methanol-chloroform precipitation. **B**. 1 vs 2 hours run of *E. Coli* proteins after acetone precipitation. **C**. 1 vs 2 hours run of human proteins after methanol-chloroform precipitation. **D**. 1 vs 2 hours run of human proteins after acetone precipitation

Methanol-chloroform precipitation resulted in 12 identified *E. Coli* proteins for both 1 and 2 hour run gradients (**Fig. 4A**). By contrast, after acetone precipitation, 13 proteins were identified in the 2-hour run, compared to 11 proteins after 1-hour run (**Fig. 4B**). With no dramatic differences at the protein level, we analysed these data at the peptide level. When examining the distribution of the peptide count following methanol-chloroform precipitation, only 2 proteins were identified with the same amount of peptides in both runs, 5 proteins had more peptides in the 2-hour run, 1 protein had more peptides in the 1-hour run, 2 proteins were present only in the 2-hour run and 4 proteins were present only in the 1-hour run (**Fig. 5A-D**). After acetone precipitation, 3 proteins were identified with the same amount of peptides in both runs, 4 proteins had more peptides in the 2-hour run, 2 proteins had more peptides in the 1-hour run, 4 proteins were present only in the 2-hour run and 2 proteins were present only in the 1-hour run (Fig. 5A-D). Overall, these data suggest that a 2-hour gradient typically identifies slightly more proteins and peptides.

**Figure 5:**
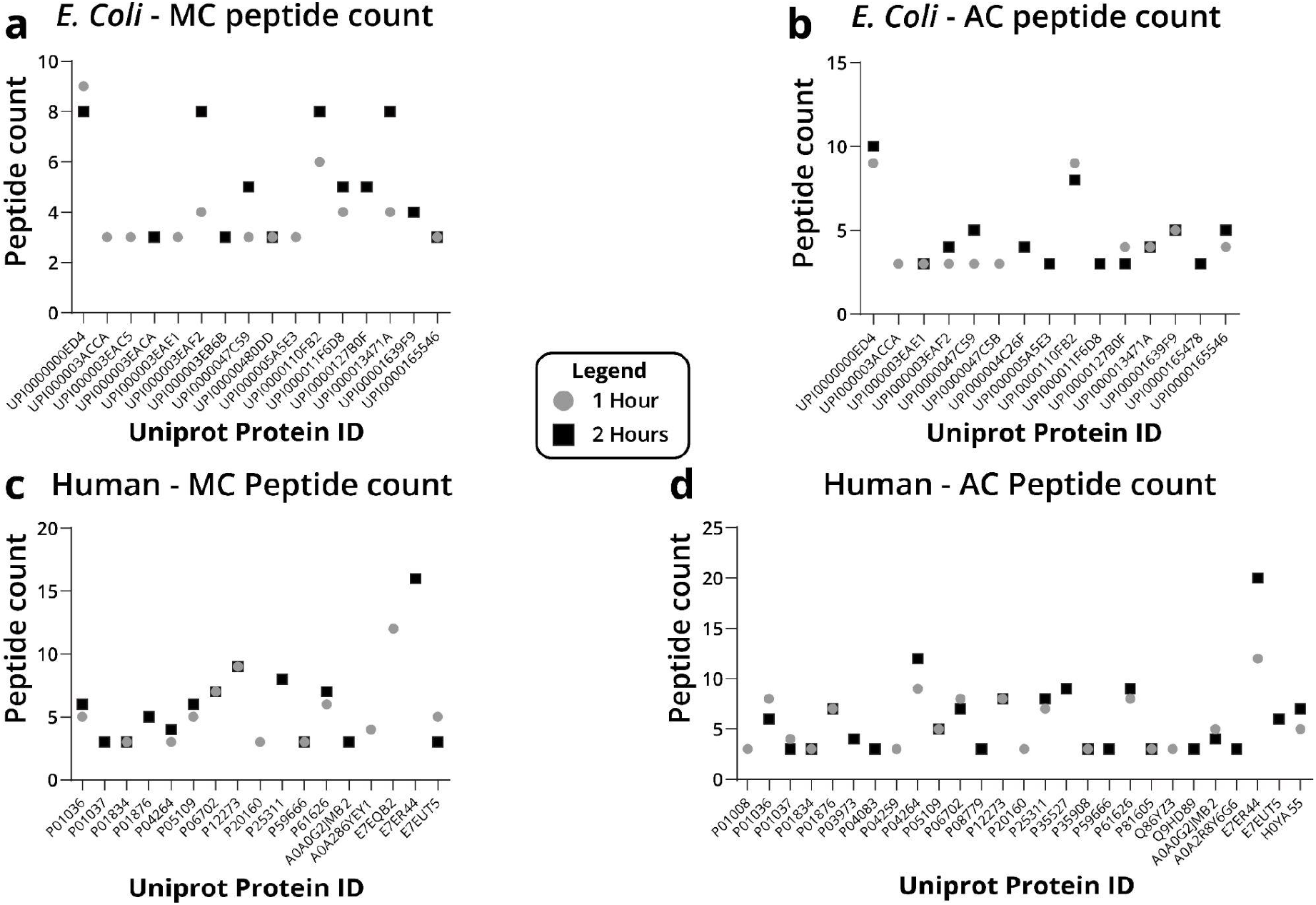
Number of peptides identified for each protein identified by 1 and 2 hour MS run and their Uniprot ID.. **A**. Peptide count of *E. Coli* proteins after methanol-chloroform precipitation. **B** Peptide count of *E. Coli* proteins after acetone precipitation. **C**. Peptide count of human proteins after methanol-chloroform precipitation. **D**. Peptide count of human proteins after acetone precipitation.

**Figure 6:**
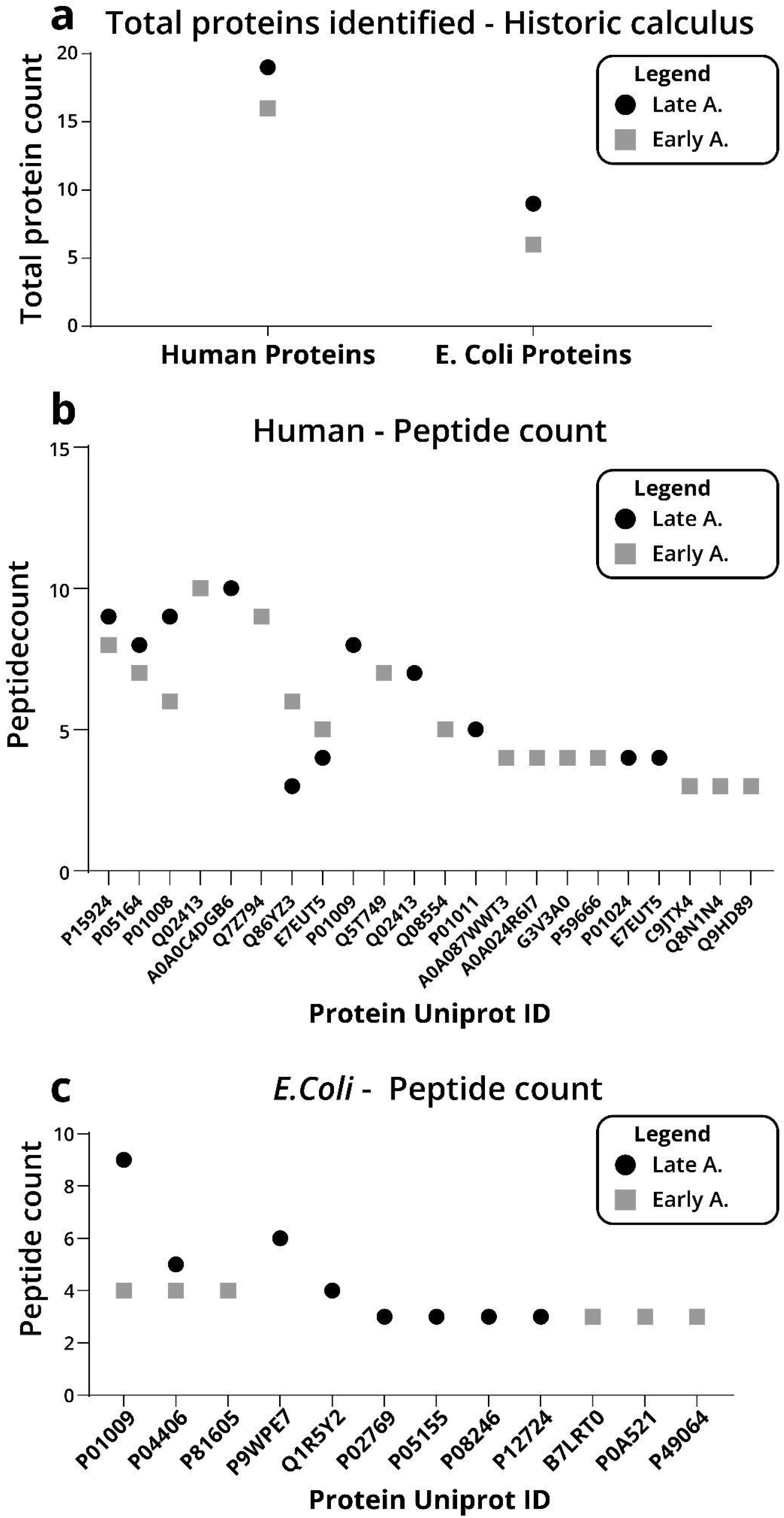
Number of peptides identified for each protein identified by 2-hour MS run. **A**. protein count of human proteins from historic dental calculus. **B** Peptide count of human proteins from historic dental calculus. **C**. Peptide count of *E. Coli* proteins from historic dental calculus.

Human proteins are known to be abundant in the saliva and gingival crevicular fluid trapped within the dental calculus ^7^. Thus, calibrating their analysis will help to identify proteins of lower abundance. In this regard, a 1-hour following methanol-chloroform precipitation identified 12 proteins, compared to 14 proteins identified in the 2-hour run (**Fig. 4C**). When comparing proteins identified after acetone precipitation, 23 proteins were identified in the 2-hours run compared to 19 in the 1-hour run (Figure 4D). In agreement with the *E. coli* data, a 2-hour gradient was likely to identify more proteins, at the expense of longer run times.

### Archaeological dental calculus analysis

To demonstrate the applicability of the established protocol to identify proteins in archaeological samples, we performed proof-of-principle experiments on two samples that were retrieved from archaeological excavations, one was dated to the 17the century (LateA) and the other was dated to the 8th century (EarlyA). In the Early Arab sample, we identified 16 human proteins and 6 *E. Coli* proteins (**Fig. 5A**). The dental calculus from the Late Arab specimen, we identified 19 human proteins and 9 *E. Coli* proteins (**Fig. 5B, C**).

## Discussion

Here, we present a protocol for proteomic analysis of modern and historic dental calculus samples. By carefully calibrating protein extraction and digestion, we were able to identify human and common bacterial proteins from modern and ancient samples. The data files generated can be used to search against databases of commonly consumed plants and animals, as well as against the oral microbiome.

A broad range of protocols to decalcify and prepare both fresh and ancient calculus for further evaluation has been previously presented. Calcification methods vary from the use of EDTA ^7,19^ to various acids at different concentrations, including nitric acid ^20^, HCl ^14,21,22^ and acetic acid ^8,17,19^, with ^20,21^ or without manual grinding, which also varies in duration and temperature of calculus denaturation. We assessed different concentrations of acetic acid and decalcification time required to extract proteins from the calcified dental calculus. Our results suggest that using higher concentration of acid followed by mechanical crushing and ultrasonic disturbance is the optimal way to extract proteins from calcified calculus.

While methanol-chloroform precipitation yielded more protein, we found that acetone precipitation provided a higher number of identified proteins and peptides. This may be the result of better retention of a large number of proteins.

Sample size is a common limiting factor in ancient calculus. We have shown that using proteomics analysis, proteins can be identified from a relatively small amount of calculus (∼10-15 mg). These amounts can be further down-scaled, in order to allow proteome analysis of ancient calculus samples, but this will likely result in a decrease in the number of proteins identified.

Ample valuable data regarding the oral microbiome, ingested food and oral and physical health can be gathered from the proteins extracted from dental calculus. Our results establish a valuable and effective protocol for gaining a considerable amount of data from a limited source of calculus. Dental calculus proteomics can be used in combination with other anthropological and archaeological research approaches aimed to increase our understanding of the biohistory of archaic populations.

## Disclaimer

The authors stated that there was no conflict of interest in conducting this research.

## Data availability

Data generated was submitted to PRIDE - Proteomics Identification Database.

## Acknowledgments

We thank the Smoler National Proteomics Centre at the Technion, Israel, for operation of the mass-spectrometer and technical assistance in sample preparation. We thank the Dan David Center for Human Evolution and Biohistory Research, Tel Aviv University, for providing the archaeological specimens. We thank the Bar and Sarig labs members for critical reading of the manuscript, useful suggestions and comments. We thank the Israeli Science Foundation (D.B. 632/20 and R.S 1339/19) for funding.

## References

1. White, D. J. Dental calculus: recent insights into occurrence, formation, prevention, removal and oral health effects of supragingival and subgingival deposits. Eur. J. Oral Sci. (1997).

2. Parfitt, G. J. A survey of the oral health of Navajo Indian children. Arch. Oral Biol. 1, 193–205 (1960).

3. Jin, Y. & Yip, H.-K. Supragingival calculus: formation and control. Crit. Rev. Oral Biol. Med. 13, 426–441 (2002).

4. Friskopp, J. & Isacsson, G. A quantitative microradiographic study of mineral content of supragingival and subgingival dental calculus. Scand. J. Dent. Res. 92, 25–32 (1984).

5. Fagernäs, Z. et al. A unified protocol for simultaneous extraction of DNA and proteins from archaeological dental calculus. J. Archaeol. Sci. 118, 105135 (2020).

6. Ziesemer, K. A. & Ramos-Madrigal, J. The efficacy of whole human genome capture on ancient dental calculus and dentin. American journal of (2019).

7. Velsko, I. M. et al. Microbial differences between dental plaque and historic dental calculus are related to oral biofilm maturation stage. Microbiome 7, 102 (2019).

8. Hendy, J. et al. Proteomic evidence of dietary sources in ancient dental calculus. Proc. Biol. Sci. 285, (2018).

9. Gismondi, A. et al. Dental calculus reveals diet habits and medicinal plant use in the Early Medieval Italian population of Colonna. Journal of Archaeological Science: Reports 20, 556–564 (2018).

10. Radini, A., Tromp, M., Beach, A., Tong, E. & Speller, C. Medieval women’s early involvement in manuscript production suggested by lapis lazuli identification in dental calculus. Advances (2019).

11. Molnar, S. Human tooth wear, tooth function and cultural variability. Am. J. Phys. Anthropol. 34, 175–189 (1971).

12. Henry, A. G., Brooks, A. S. & Piperno, D. R. Microfossils in calculus demonstrate consumption of plants and cooked foods in Neanderthal diets (Shanidar III, Iraq; Spy I and II, Belgium). Proc. Natl. Acad. Sci. U. S. A. 108, 486–491 (2011).

13. Geber, J. et al. Relief food subsistence revealed by microparticle and proteomic analyses of dental calculus from victims of the Great Irish Famine. Proc. Natl. Acad. Sci. U. S. A. 116, 19380–19385 (2019).

14. Warinner, C. et al. Pathogens and host immunity in the ancient human oral cavity. Nat. Genet. 46, 336–344 (2014).

15. Akcalı, A. & Lang, N. P. Dental calculus: the calcified biofilm and its role in disease development. Periodontol. 2000 76, 109–115 (2018).

16. Kenward, H. & Hall, A. Environmental Archaeology Unit Reports. https://www.york.ac.uk/inst/chumpal/EAU-reps/eaureps-web.htm.

17. Jersie-Christensen, R. R. et al. Quantitative metaproteomics of medieval dental calculus reveals individual oral health status. Nat. Commun. 9, 4744 (2018).

18. Hardy, K. et al. Starch granules, dental calculus and new perspectives on ancient diet. J. Archaeol. Sci. 36, 248–255 (2009).

19. Bleasdale, M., Richter, K. K., Janzen, A. & Brown, S. Ancient proteins provide evidence of dairy consumption in eastern Africa. Nature (2021).

20. Zhang, B., Tan, X. & Zhang, K. Cadmium Profiles in Dental Calculus: a Cross-Sectional Population-Based Study in Hunan Province of China. Biol. Trace Elem. Res. 185, 63–70 (2018).

21. Yaprak, E. et al. High levels of heavy metal accumulation in dental calculus of smokers: a pilot inductively coupled plasma mass spectrometry study. J. Periodontal Res. 52, 83–88 (2017).

22. Gismondi, A. et al. A multidisciplinary approach for investigating dietary and medicinal habits of the Medieval population of Santa Severa (7th-15th centuries, Rome, Italy). PLoS One 15, e0227433 (2020).

