## Supplementary File 1 for "The hidden secrets of the dental calculus: Calibration of a mass spectrometry protocol for dental calculus protein analysis"

**Decalcification:**

1. Wash sample with 500ul PBS for 15 min in a 1.7ml tube in a 3D orbital shaker
2. Centrifuge for 10 min at 10000 RCF and remove buffer.
3. Repeat the wash and centrifuge (1-2) 3 times.

**Demineralization**

4. Add 500ul of 25% acetic acid in DDW, leave overnight in a 3D orbital shaker.
5. Crush the calculus with micropestels
6. Centrifuge for 10 min, at 4C and 10,000 RCF.
7. Remove the acetic acid, add 300ul PBS and vortex (up and down)
8. Repeat the washes 4 times.
9. After the last centrifuge replace the PBS with 150ul lysis buffer (pH 7.5: 2M Guanidine Hydrochloride (Merck), 10mM Chloroacetamide (Bar Naor - Israel) and 5mM HEPES (Merck)
10. Sonication for 25 min at 40% (Sonic Ruptor 400, Omni)
11. After sonication, spin down for 30 min at 4C at 21,000 RCF
12. Separate the pellet from the supernatant, keep supernatant.

**Precipitation- choose preffered method:**

**Aceton (recommended):**

13. Add 600 ul of cold (-20C) acetone (equal to 4 times sample volume) and vortex thoroughly.
14. Put the sample in -80C for 60 min.
15. Centrifuge at 15,000 RCF for 10 min at 4C, discard or acetone while keeping the pellet.
16. Using a speedvac or a desiccator, completely remove acetone remains.
17. Resuspend with 100ul of 100mM HEPES.

**Methanol- chloroform (alternative):**

18. For a sample volume of 150ul add 600ul methanol and vortex.
19. Add equal amounts of chloroform to the original sample (150 ul) and vortex.
20. Add 400ul of double distilled water and vortex thoroughly, until the mixture is cloudy with precipitation.
21. Centrifuge at 14,000 RCF for 1 min.
22. Remove the top layer without disturbing the circular flake of protein.
23. Add methanol equal to 4 time of sample volume and vortex.
24. Centrifuge for 5 min at 20,000 RCF.
25. Discard excess methanol.
26. Lufilize the sample overnight.
27. Resuspend with 100ul of 100mM HEPES.

**Reduction:**

28. Add Dithiothreitol to a final concentration of 10mM
29. Incubate in shaker for 60 min at 55C and 200rpm

**Alkylation:**

30. Immediately before use, add 100 mM HEPES to Iodoacetamide to a final concentration of 18.75mM Iodiacetamide
31. Incubate for 30 min at room temperature covered with tinfoil**.**

**Trypsinization:**

32. Add 10nG mass spectrometry grade trypsin (2 uL of 5 nG / uL stock)
33. Incubate overnight at 37 C and 200rpm
34. After overnight digestion, stop trypsinization by adding 1ul of Trifluoroacetic acid (TFA 0.1%)

**Solid Phase Extraction Stage Tips for detergent removal:**

**C18 Empore™ Stage - Tip:**

35. Use 2 layers of C18 Empore™ SPE disks,punched and inserted into 200ul tips.
36. Activate with 100ul MeOH
37. Centrifuge for 2 min at 1500 RCF and discard excess fluids.
38. Activate and cleaning from residual peptides with 100ul buffer B (80% acetonitrile,0.1% TFA)
39. Centrifuge for 2 min at 1500 RCF and discard excess fluids.
40. Return to hydrophilic state with 100ul buffer A (0.1%TFA)
41. Load up to 6 ug protein
42. Centrifuge for 2 min at 1500 RCF and discard excess fluids.
43. Return to hydrophilic state with 100ul buffer A (0.1%TFA)
44. Repeat the previous step
45. Manually elute with 60 ul of buffer B.
46. Put the tube with the elution under vacuum.
47. Can be stored at room temperature.

**SCX strong cation exchange Empore™ Stage Tip:**

48. Use 2 layers of C18 Empore™ SPE disks.
49. Wash the layers with 400ul of TFA 0.1%
50. Resuspend the sample with 100ul TFA 0.1%
51. Load the sample and pass it through manually
52. Wash with 200ul TFA 0.1%
53. Elution with 100ul methanol 30% ammonium hydroxide 5%
54. Put under vacuum over night.

**Liquid Chromatography and mass spectrometry:**

55. Resolve peptides by reverse phase chromatography on 0.075 X 180 mm fused silica capillaries (J&W) packed with Reprosil reverse phase material (Dr Maisch GmbH, Germany).
56. Elute peptides with a 60 or 120 minutes linear gradient of 5% to 28% 15 minutes gradient of 28 to 95% and 25 minutes at 95% acetonitrile with 0.1% formic acid in water at flow rates of 0.15 µl/min.
57. Perform mass spectrometry by Q Exactive HF mass spectrometer (Thermo) in a positive mode using repetitively full MS scan followed by collision induced dissociation of the 18 most dominant ions (>1 charges) selected from the first MS scan.
58. Perform a dynamic exclusion list with exclusion duration of 20 seconds.
**59.**
